# Distinct, opposing functions for CFIm59 and CFIm68 in mRNA alternative polyadenylation of *Pten* and in the PI3K/Akt signalling cascade

**DOI:** 10.1101/2021.09.09.459613

**Authors:** Hsin-Wei Tseng, Anthony Mota-Sydor, Rania Leventis, Ivan Topisirovic, Thomas F. Duchaine

## Abstract

The precise maintenance of PTEN dosage is crucial for tumor suppression across a wide variety of cancers. Post-transcriptional regulation of *Pten* heavily relies on regulatory elements encoded by its 3’UTR. We previously reported the important diversity of 3’UTR isoforms of *Pten* mRNAs produced through alternative polyadenylation (APA). Here, we reveal the direct regulation of *Pten* APA by the mammalian cleavage factor I (CFIm) complex, which in turn contributes to PTEN protein dosage. CFIm consists of the UGUA-binding CFIm25 and APA regulatory subunits CFIm59 or CFIm68. Deep sequencing analyses of perturbed (KO and KD) cell lines uncovered the differential regulation of *Pten* APA by CFIm59 and CFIm68 and further revealed that their divergent functions have widespread impact for APA in transcriptomes. Differentially regulated genes include numerous factors within the phosphoinositide 3-kinase (PI3K)/protein kinase B (Akt) signalling pathway that PTEN counter-regulates. We further reveal a stratification of APA dysregulation among a subset of *PTEN-driven* cancers, with recurrent alterations among PI3K/Akt pathway genes regulated by CFIm. Our results refine the transcriptome selectivity of the CFIm complex in APA regulation, and the breadth of its impact in *PTEN*-driven cancers.

## INTRODUCTION

Alternative polyadenylation (APA) of messenger RNAs (mRNAs) has emerged as a fundamental layer of gene regulation and is thought to contribute to the tuning of at least 70% of all mammalian mRNA-coding genes (1,2). APA greatly expands the transcript diversity through utilization of multiple polyadenylation signals (PAS) that lead to expression of mRNA isoforms with varying 3’ untranslated region (3’UTR) lengths. PAS that are proximal to the coding sequence (CDS) give rise to shorter 3’UTR isoforms and can exclude *cis*- regulatory elements that otherwise would be encoded in longer 3’UTR isoforms. This reorganization severs the 3’UTR from its connections with trans-acting factors such as RNA-binding proteins (RBPs), microRNAs (miRNAs), the associated RISC, and their effector machineries, and further affects 3’UTR folding structures. As such, APAs can have profound impact on mRNA translation, stability, localization, and functions that are often closely linked to the molecular and transcriptional makeup of cell identities (3). APA analysis of single cell RNA-seq datasets also robustly delineated cell subpopulations in tumors and through spermatogenesis (4). Aberrant regulation of APA has been linked with human disease, such as through oncomir targeting (5), and is prominent in cancer wherein it is often associated with marked global shortening of 3’UTRs (6–8).

*P*hosphatase and *TEN*sin homolog (*PTEN*) is one of the most frequently inactivated tumor suppressor genes in cancer (9). Its activity antagonizes the PI3K/Akt signalling pathway, thereby protecting cells against unchecked cell proliferation, invasion, or evasion of programmed cell death (10). Its expression must be tightly regulated as even a partial loss of *PTEN* expression can increase the likelihood of tumor formation (11,12). Incidentally, *PTEN* mRNAs undergo extensive APA regulation, with up to 6 distinct lengths of 3’UTR observable in mouse cell lines, and even more in human cells. We had previously reported that longer isoforms exhibit greater stability and contribute to the bulk of protein expression (13). Resolving how APA itself is regulated and how PTEN dosage is controlled through APA are important to understand gene dysregulation in cancer. The mammalian 3’ end processing apparatus is composed of 16 ‘core’ proteins associated into the CPSF, CstF, CFIm and CFIIm complexes and a poly(A) polymerase (14,15) and molecular architectures of some of the cleavage and polyadenylation machinery has been detailed (16). The location of pre-mRNA 3’ end is mainly defined by consensus poly(A) signals defined by the canonical sequence AAUAAA (17), and the cleavage site ~15 nt downstream, where polyadenylation occurs. Potency and selectivity of PAS are thought to rely on surrounding *cis*- regulatory elements and input from a number of global APA regulators that were only recently identified (3,18). Of particular interest for this function is the mammalian cleavage factor I (CFIm), which targets the UGUA motif in the proximity of PAS. At the molecular level, functional UGUA sites can potentiate cleavage through nearby PAS, but is not essential for cleavage (19). Biochemical and structural studies described CFIm as a heterotetrameric complex consisting of a CFIm25 dimer, which directly recognizes the UGUA sequence, and a dimer of CFIm59 or CFIm68 (20–22). CFIm subunit dysregulation has been associated with glioblastoma (GBM) aggressiveness and with neuropsychiatric diseases (23,24). Curiously, it has been reported that depletion of CFIm25 or CFIm68, but not CFIm59, leads to global shortening of 3’UTRs (19,25). CFIm59 function is thought to be partly redundant with CFIm68 as it partially rescues APA shift caused by CFIm68 depletion (19,20), but the specific functions of CFIm59 in APA regulation remain elusive.

Here we identified CFIm as a direct regulator of *PTEN* APA with significant influences over its mRNA and protein expression. We uncovered distinct and mostly opposing effects for CFIm68 and CFIm59 depletion on APA, not only for *PTEN* but transcriptome-wide, including for several cancer genes. Collectively, our results provide a view of the precise roles for CFIm subunits in APA regulation of PTEN dosage, a broad impact on APA in the PI3K/Akt cascade, and on the overall transcriptome.

## MATERIAL AND METHODS

### Cell culture and transfection

NIH3T3 cells (ATCC) were cultured in Dulbecco’s modified Eagle medium (DMEM) with 10% calf serum (Cytiva). HEK293T cells kindly shared by Dr. Alan Engelman and Dr. Yonsheng Shi labs were cultured in DMEM supplemented with 10% fetal bovine serum (FBS) (Wisent). All cell lines were grown in a humidified incubator at 37 °C with 5% CO2. Knockdown in NIH3T3 was performed by reverse transfection of siRNAs using Lipofectamine RNAiMAX (ThermoFisher) with a final concentration of 10 μM for 72 hours. The following siRNAs were used: negative ctrl (Qiagen 1027281), CFIm25-si1 (Qiagen SI04414732), CFIm25-si2 (Qiagen SI04414739), CFIm59-si1 (Qiagen SI00858655), CFIm59-si2 (forward: 5’-GUCCUCAUCUCCUCUCUUATT-3’, reverse: 5’-UAAGAGAGGAGAUGAGGACTT-3’), CFIm68-si1 (Qiagen SI00958741), and CFIm68-si2 (Qiagen SI00958748).

### RNA isolation and Northern blots

Total RNA from mammalian cell lines was extracted with either the miRNEasy mini kit (Qiagen) or the Monarch total RNA miniprep kit (NEB). Depending on the level of expression, 1-4 μg of total RNA was loaded on 1% agarose-glyoxal gel prepared with NorthernMax-Gly gel prep running buffer (Ambion) and transferred to nylon membranes (BrightStar-Plus, Invitrogen). Probes were synthesized using DECAprime II (Invitrogen) with α-^32^P dATP, or MEGAscript T3 kit (Invitrogen) with α-^32^P UTP. Hybridization was performed overnight at 42 or 68 °C in ULTRAHyb hybridization buffer (Ambion). Membranes were then washed and exposed overnight onto imaging plates and imaged on Typhoon phosphorimager (GE). Primers used to amplify both human and mouse *Pten* probes were (TDO1774) 5’-ACCAGGACCAGAGGAAACCT-3’ and (TDO1775) 5’-GAATGCTGATCTTCATCAAAAGG-3’.

### *Pten* 3’UTR isoform-specific qRT-PCR

The protocol used was developed in (13). Briefly, cDNA was generated from 500 μg of total RNA using an oligo-dT12 anchor primer (TDO679, see Supplementary Table 1) with SuperScript III reverse transcriptase (Invitrogen). To quantify isoforms 300 and 3.3k, first-round amplification (PCR1) was performed with 1:4 diluted reverse transcription (RT) reaction using isoform specific primers (Supplementary Table 1) with a short extension time. The following round of real-time PCR (qPCR2) was performed with 1:1,000 or 1:100,000 diluted PCR1 product using SsoAdvanced Universal SYBR green supermix (Bio-Rad). To quantify the *Pten* 5-6k isoform, total *Pten* (targeting ORF), and Hprt1 internal control, qPCR was performed directly on the 1:4 diluted RT reaction without the PCR1 step. Relative expressions were quantified using the ΔΔCt method and normalized to Hprt1.

### Reporter constructs

Mouse *Pten* sequences were cloned into the pcDNA4/TO plasmid, including the coding sequence and the 6.1k 3’UTR from the genomic sequence. Mutagenesis was performed on each targeted UGUA using all-around PCR with non-overlapping primers. Primers used for mutations are detailed in Supplementary Table 1.

### Western blotting

Total cell lysates were prepared using lysis buffer (50 mM Tris-HCl pH 7.4, 150 mM NaCl, 1% IGEPAL CA-630, 1% sodium deoxycholate, 0.1% sodium dodecyl sulphate, and 1 mM ethylenediaminetetraacetic acid) supplemented with protease and phosphatase inhibitors (Sigma and Roche). Proteins were immobilized on PVDF membranes (Millipore) and probed with the following antibodies diluted in the Odyssey blocking buffer (Li-COR): anti-CFIm68 (Bethyl A301-358A-T, 1:1,000), anti-CFIm25 (Proteintech 10322-1-AP, 1:250), anti-CFIm59 (Sigma HPA041094, 1:500), anti-β-actin (CST 8H10D10, 1:5,000), anti-PTEN (CST 9552, 1:1,000). Secondary IR dye antibodies against mouse or rabbit (Li-COR) were used at 1:10,000. Blots were scanned using the Li-COR imaging system and quantified with the Image Studio Lite suite.

### 3’UTR-seq and data processing

Using 500 ng of total RNA prepared as described, 3’UTR libraries were generated using QuantSeq 3’ mRNA library prep kit REV (Lexogen) following the manufacturer’s instructions. Quality control and sequencing of the libraries were performed by the IRIC genomics platform (Université de Montréal) using NextSeq500 SR75 to obtain approximately 20 million reads per sample. Raw reads were preprocessed to remove any adapter sequences using Trimmomatic v0.36 and mapped to mm10 genome with STAR v2.7.2b. Subsequent identification and quantification of PAS peaks were performed according to (26). Briefly, reads from all samples were merged for peak calling using CLIPanalyze (https://bitbucket.org/leslielab/clipanalyze). Internal priming events were then removed along with low-usage events to compile a curated list of valid APA sites. This list was then referenced to count APA events in each sample using featureCounts (27). Differential expression of each APA event between conditions was performed through repurposing the DEXSeq package (28) originally developed for the analysis of differential exon expression.

For RED score analyses, genes with only one identified PAS were first filtered out. The top two expressing PAS across all samples of each remaining gene were identified and defined as the proximal or distal site depending on their relative position. Responsive genes are defined as |RED| ≥ log_2_(1.5).

### Statistical analyses

Plotted quantification of western blots, northern blots and qRT-PCR are all presented as mean ± standard deviation. Statistical tests were performed using one-way Analysis Of Variance (ANOVA) with post-hoc Dunnett’s test between all treatment and control groups. All plots were generated using the gglot2 package (29) in R.

## RESULTS

### CFIm regulates *Pten* mRNA and protein expression

To determine whether PTEN protein dosage is regulated through alternative polyadenylation (APA) of its transcripts, we first investigated the role of CFIm, a known APA regulator complex. To this end, we knocked down (KD) each member of the CFIm complex (CFIm25, CFIm59, and CFIm68) in mouse fibroblast cell line NIH3T3 with two different siRNAs, each of which achieving at least 70% KD, and probed for PTEN protein expression by western blot (WB). Depletion of individual CFIm members resulted in significant upregulation of PTEN protein, with comparable increases ranging from 1.3 to 1.8-fold across CFIm members (Figure 1A, 1B). KD of subunits also consistently led to the upregulation of *Pten* mRNAs. qRT-PCR on *Pten* open reading frame (ORF) detected a ~2.5-fold mRNA upregulation upon CFIm25 KD, between 1.5 to 2.7-fold for CFIm59 KD, and between 2.2 to 4-fold for CFIm68 KD (Figure 1C). Variation across replicates and the apparently more limited changes in protein expression are consistent with the known multi-layered mechanisms controlling PTEN dosage which may be partially buffering the impact of CFIm KD (see Discussion). Interestingly, we noticed an inter-dependence of CFIm subunits for their expression. Depletion of CFIm25 led to a reduction of both CFIm59 and CFIm68 proteins, whereas depletion of CFIm59 or CFIm68 led to a reduction of the CFIm25 subunit. However, KD of CFIm59 and CFIm68 expression did not affect one another (Figure 1D). Coordinated expression of the complex subunits is likely co- or post-translational as mRNA-seq analyses from libraries derived from KD (Supplementary Figure 1, also see below) could not explain this down-regulation. These results demonstrate the functional relevance of CFIm as an important regulator of PTEN dosage and suggest an impact for this complex on *Pten* mRNA APA.

**Figure 1.**
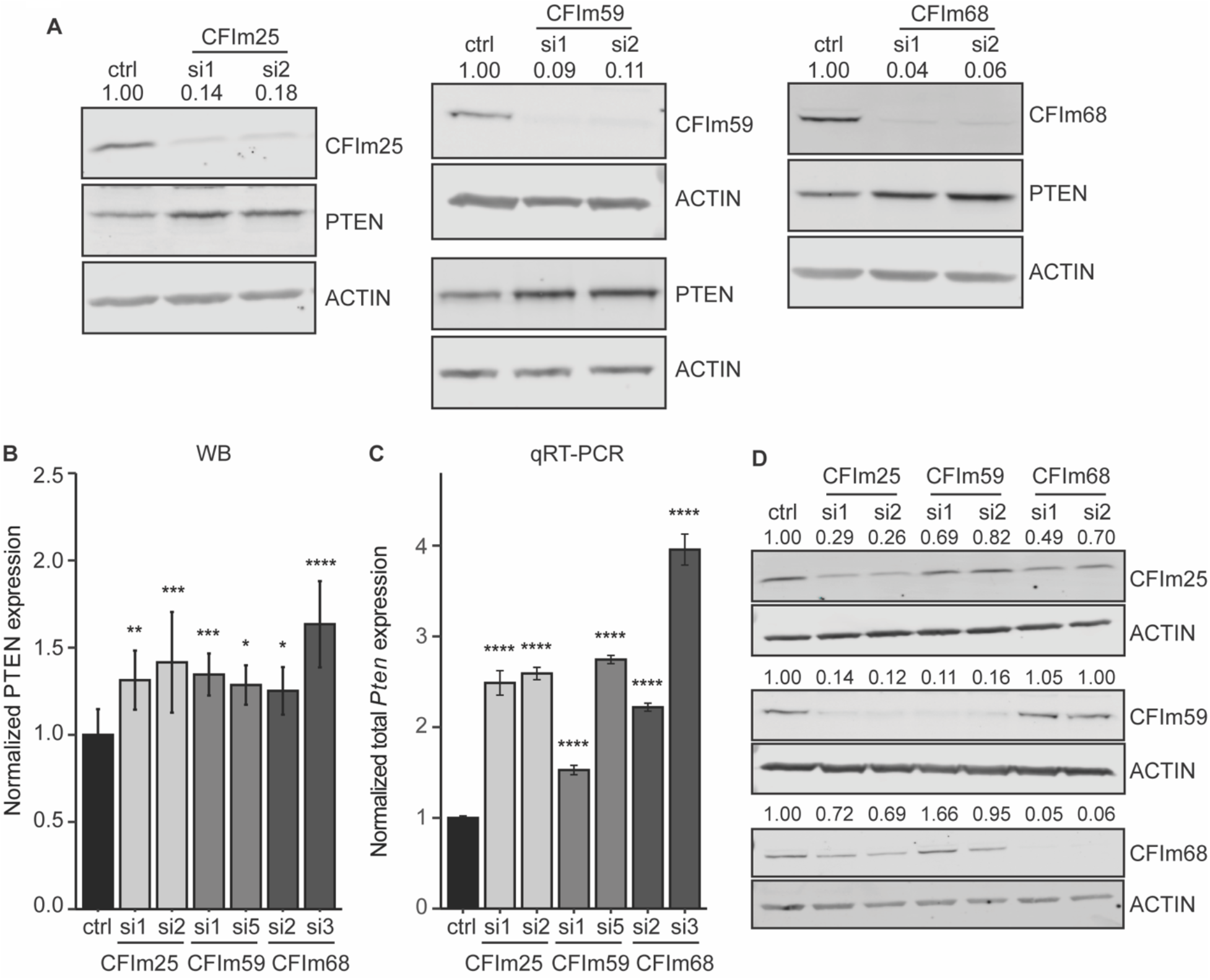
CFIm regulates *Pten* mRNA and protein expression. (A) Representative western blots of CFIm25, −59, and −68 KD and the corresponding change in PTEN level. Two different siRNAs were used for each KD, and the efficiency was quantified. Actin was used as a loading control. (B) Quantification of normalized PTEN protein expression upon CFIm KD. (C) Quantification of normalized total *Pten* mRNA expression upon CFIm KD assayed through qRT-PCR and normalized to *Hprt*. (D) Western blotting of CFIm25, −59, and −68 showing codependent protein expression. All error bars indicate mean ± standard deviation across at least three biological replicates. P-values were calculated with one-way ANOVA followed by Dunnett’s test (\**P* ≤ 0.05, \*\**P* ≤ 0.01, \*\*\**P* ≤ 0.001 and \*\*\*\**P* ≤ 0.0001).

### CFIm controls *Pten* mRNA 3’UTR isoform expression

To determine how KD of CFIm subunits leads to *Pten* upregulation, we investigated the impact of their disruption on *Pten* mRNA APA. Using 3’UTR-seq (Lexogen), we observed multiple *Pten* 3’UTR isoforms that we had previously detected by 3’ RACE and unambiguously mapped (13). The two major *Pten* 3’UTR isoforms are defined by a proximal PAS located at 279 nt, and a cluster of 4 PAS located at 3255, 3276, 3312, and 3351 nt downstream of the stop codon, respectively. The 4 PAS near the 3.3k nt mark are clustered in a 102-nt stretch of the 3’UTR, their output is practically indistinguishable and will thus be considered together here. The predominant isoforms will be referred to as APA 300 nt and APA 3.3k. Lesser expressed isoforms between 5.4k and 6.1k nt were also consistently observed and will altogether be referred to as APA 5-6k nt.

KD of CFIm25 consistently shifted APA usage towards the proximal 300 nt isoform, at the expense of the longer 3.3k and 5-6k nt isoforms (Figure 2A). KD of CFIm68 resulted in an even more extensive APA shift towards the 300 nt isoform (Figure 2A). Intriguingly, knockdown of CFIm59 triggered a distinct and opposite shift, with longer isoforms (3.3k and 5-6k nt) becoming significantly favored relative to the proximal 300 nt isoform (Figure 2A). These results were further corroborated using northern blot on CFIm knockdown samples. The proximal (300 nt)-to-distal (3.3k and 5-6k nt) isoforms ratio increased in CFIm25 and CFIm68 KD and decreased in CFIm59 KD (Figure 2B). Lastly, we further quantified *Pten* 3’UTR APA upon CFIm depletion using isoform-specific qRT-PCR (13). For CFIm25 and CFIm68 KD, the 300 nt 3’UTR isoform was increased 5-fold and 6-10–fold, respectively (Figure 2C), and this was accompanied by a significant reduction of distal isoforms. In contrast, CFIm59 KD led to the up-regulation of the 3.3k isoform by 1.8 to 3-fold, and up-regulation of the 5-6k nt isoforms by 2 to 4-fold, with no detected changes in the 300 nt isoform using this assay (Figure 2C). Altogether, this data provides compelling evidence of *Pten* APA regulation by all CFIm subunits and indicate that CFIm59 may exert a distinct and opposing function to CFIm68. We further noticed from both northern and qRT-PCR results, that APA redistribution was accompanied by an increase in overall *Pten* mRNA abundance (Figure 1C). In the case of CFIm68 KD, this was mainly driven by over-expression of the proximal 300 nt isoform, while CFIm59 KD instead led to a milder but significant increase in distal 3.3kb and 5-6kb isoforms. These results suggest that the increase in PTEN protein expression upon CFIm59 and CFIm68 KD can at least partly be accounted for by the translation of distinct *Pten* mRNA isoforms (see Discussion).

**Figure 2.**
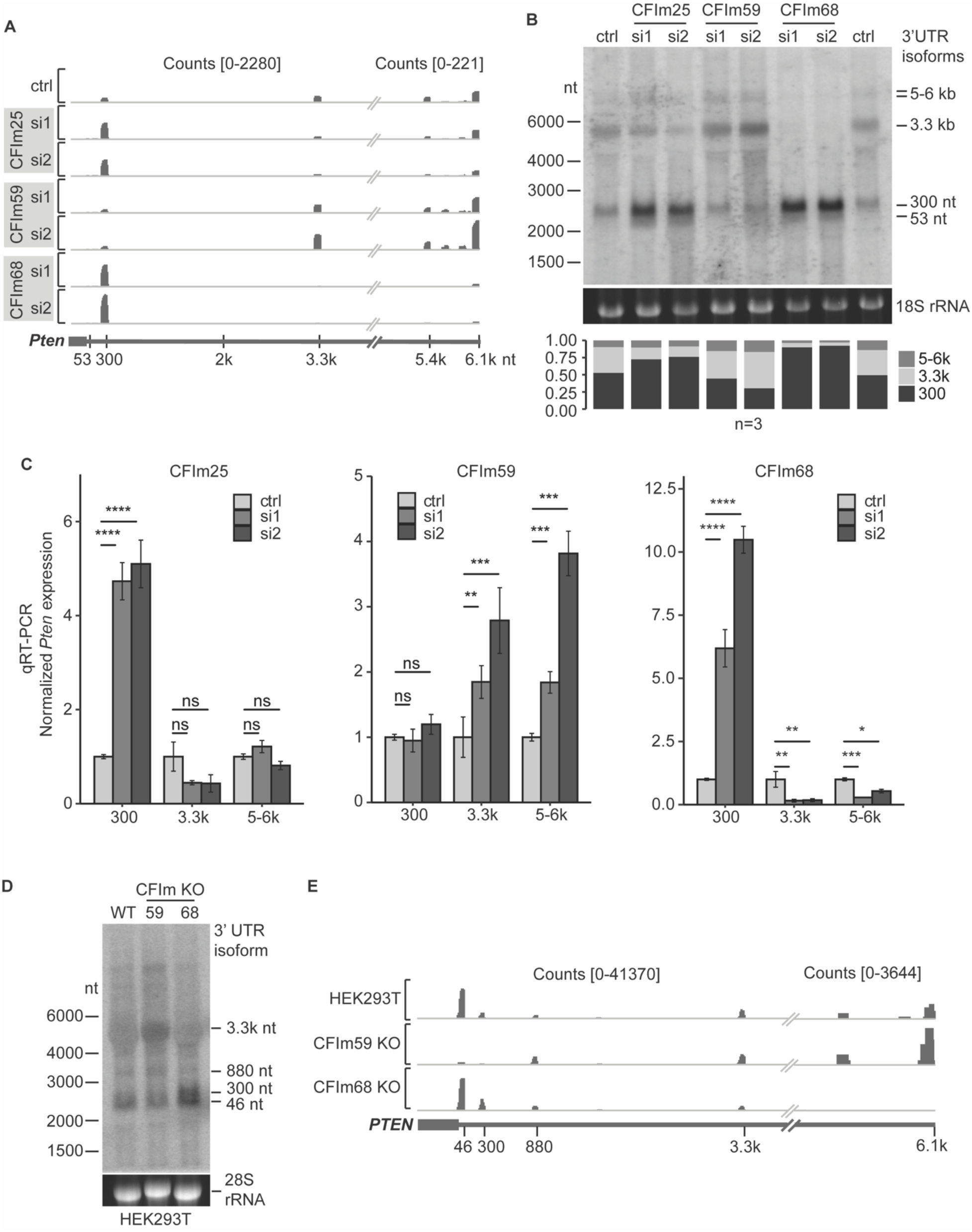
CFIm controls *Pten* mRNA 3’UTR isoform expression. (A) Representative *Pten* tracks of 3’UTR-seq upon CFIm25, −59, and −68 KD in NIH3T3. Schematic of *Pten* and 3’UTR isoforms previously captured by 3’ RACE (13) is shown on the bottom. Each track is separated into two parts with different scales indicated as count ranges on top to show all major isoforms. (B) Quantitative northern blotting of endogenous *Pten* upon CFIm KD in NIH3T3. Corresponding 3’UTR isoforms are indicated on the right with the size markers on the left. Bar graph on the bottom tabulates the average relative composition of the 300 nt, 3.3k and 5-6k isoforms across three biological replicates. (C) *Pten* 3’UTR isoform-specific qRT-PCR for the 300 nt, 3.3k and 5-6k isoforms upon CFIm KD in NIH3T3. (D) Northern blotting of endogenous human *PTEN* in HEK293T and isogenic cell lines CFIm59 and −68 KO. Size markers are shown on the left and the identified 3’UTR isoforms on the right. (E) PAS-seq tracks of HEK293T and isogenic CFIm59 and −68 KO cell lines. Schematic of *PTEN* PAS appears on the bottom. The tracks are separated into two parts with different scales of counts indicated on top. All experiments were performed in three biological replicates with technical triplicates for qRT-PCR. Error bars indicated are mean ± standard deviation. P-values were calculated with one-way ANOVA followed by Dunnett’s test (\**P* ≤ 0.05, \*\**P* ≤ 0.01, \*\*\**P* ≤ 0.001, \*\*\*\**P* ≤ 0.0001, and ns not significant).

### Conservation and species-specific functions of CFIm on human *PTEN*

The 3’UTR sequences of human and mouse *Pten* share 74% sequence identity. The predominant proximal (300 nt) and distal (3.3k) PAS as well as the nearby UGUA elements are conserved in human 3’UTR-encoding sequences. The human *PTEN* 3’UTRs further encode additional frequently used PAS, including an additional proximal PAS at 46 nt and another site at 880 nt (13). We took advantage of the HEK293T isogenic cell line panel to compare the effect of CFIm on human *PTEN* 3’UTRs. Northern blot on endogenous *PTEN* mRNAs led to sensibly the same observations as with mouse sequences (Figure 2D). CFIm68 KO strongly favored use of proximal APAs, whereas CFIm59 KO significantly increased distal PAS 3’UTR isoforms in comparison with the parental cell line. These results were further corroborated by mining *PTEN* 3’UTRs from published PAS-seq datasets of CFIm59 KO and CFIm68 KO lines (19) (Figure 2E). Strikingly, APA 46nt *PTEN* 3’UTR was drastically reduced in CFIm59 KO, but its use was not significantly altered in CFIm68 KO, indicating that this human-favored proximal APA is exclusively and acutely sensitive to CFIm59 function.

Together, these results highlight the conservation of distinct, opposite roles of CFIm59 and CFIm68 in human and mouse in *PTEN* mRNA APA, and further reveal some species-specific differences in 3’UTR regulatory sequences.

### CFIm subunits and UGUA RNA elements in *Pten* APA regulation

The CFIm complex recognizes the UGUA consensus motif near PAS through its CFIm25 RNA-binding subunit (21,30). To determine the direct contribution of CFIm to *Pten* APA regulation, we identified and mutagenized UGUA sites near the two major PAS in mouse 3’UTRs. Three candidate sites surround APA 300 nt, including UGUA1 and UGUA2 at 54 nt and 45 nt upstream, and UGUA3 at 55 nt downstream (Figure 3A). Two candidate CFIm binding sites are encoded near the 3.3k PAS (UGUA4 and UGUA5), at 127 nt and 94 nt upstream of the second and most used PAS in the cluster, as determined previously by 3’RACE (13). We engineered an APA reporter by cloning the full-length mouse *Pten* ORF and 6.1 kb of its 3’UTR genomic sequence in a CMV-driven construct, and the five <u>U</u>G<u>U</u>A candidate sites were mutated to <u>A</u>G<u>A</u>A, individually or in combinations (Figure 3A). Similar mutation was previously used to incapacitate CFIm recognition and cleavage *in vitro* (30). Constructs were transfected transiently in human HEK293T cells and usage of *Pten* PAS was monitored and quantified using mouse-specific *Pten* mRNA northern blotting. The lack of signal in the empty-vector transfected cells demonstrated the specificity of the northern (Figure 3B, 3D, empty). WT reporter transfection led to expression of *Pten* 300 nt, 3.3k, and 5-6k 3’UTR isoforms, consistent with what was observed endogenously (Figure 3B). Relative use of proximal to distal PAS (300 nt/3.3k isoforms) was quantified for each UGUA mutation and normalized to WT (Figure 3C, E). Individually disrupting UGUA2 located upstream of APA 300 nt significantly shifted isoform ratio in favor of distal APA 3.3k, while mutation of the upstream UGUA1 or downstream UGUA3 alone had no significant effect. Individual mutation of either of the two UGUA that are proximal to APA 3.3k (UGUA 4 or UGUA5) did not significantly affect *Pten* APA. Double mutants with UGUA2 mutation decreased the 300/3.3k ratio to an extent comparable with the single UGUA2 mutation alone. However, triple UGUA1, 2, and 3 mutations near APA 300 nt further decreased the 300/3.3k ratio, favoring the 3.3k isoform, with contributions that appeared to be mostly additive. In contrast, combining mutations in UGUA4 and 5 greatly increased the magnitude of 300/3.3k ratio, over either UGUA 4 or UGUA5 mutations, suggesting a possible compensation between the two sites (Figure 3B, 3C). Interestingly, when all five candidate UGUA sites were mutated, the 3.3k isoform prevailed over APA 300 nt, suggesting that distal isoforms are intrinsically favored, independently of UGUA and/or CFIm. This may be influenced by cell culture conditions such as density (13) and/or input from additional, yet unidentified APA regulatory elements in *Pten* 3’UTR.

**Figure 3.**
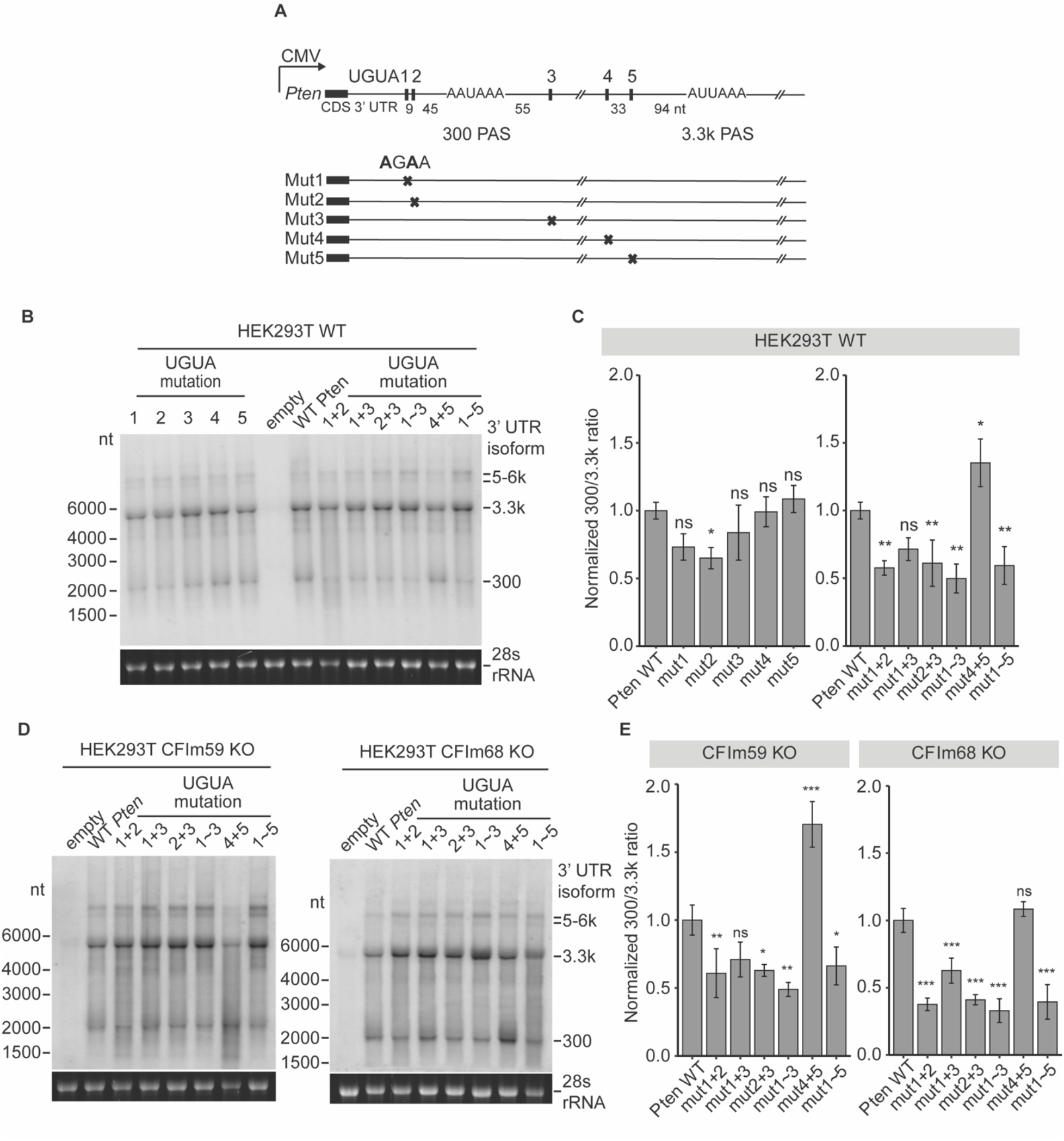
CFIm subunits and UGUA RNA elements in *Pten* APA regulation. (A) Schematic of five UGUA motifs surrounding mouse *Pten* PAS 300 nt and PAS 3.3k on a CMV-driven construct. Distance between the PAS and each UGUA is indicated. Bottom shows individual mutations introduced at each UGUA site. (B, D) Mouse *Pten*-specific northern blotting of HEK293T wildtype (B), CFIm59 KO and CFIm68 KO cells (D) transfected with *Pten* constructs. Size markers are indicated on the left and the *Pten* 3’UTR isoforms on the right. Empty: mock transfection with construct backbone only. (C, E) Quantification of (B) and (D) across three biological replicates. Ratio of 300 nt and 3.3k isoforms was calculated for each transfected sample and normalized to the ratio of wildtype (WT) *Pten*-transfected sample. Error bars are mean ± standard deviation and P-values were calculated with one-way ANOVA followed by Dunnett’s test (\**P* ≤ 0.05, \*\**P* ≤ 0.01, \*\*\**P* ≤ 0.001, and ns not significant).

To investigate the impact of CFIm subunits CFIm59 and CFIm68 on UGUA sites, we profiled reporter APA use by transfecting isogenic HEK293T lines bearing the CFIm59 or CFIm68 KO (19,31). As in the parental cell line, KO of either CFIm59 or CFIm68 significantly decreased the 300/3.3k APA ratio with combined mutations in proximal UGUA sites 1, 2, and 3 (Figure 3D, E). The combined effect of mutation in the 3 proximal UGUA elements (1, 2, 3) in CFIm59 KO cells was indistinguishable from the parental line (Figure 3C, 3E). This suggests that the shift towards distal PAS upon CFIm59 depletion does not require the proximal UGUA sites. In contrast, CFIm68 KO exacerbated the impact of mutations in proximal UGUA1-3, further enhancing the use of distal PAS over their effect in the parental line (Figure 3C right, 3E right). Furthermore, whereas mutation of distal UGUA4+5 in CFIm59 KO led to greater enhancement of proximal PAS use over the parental line, the same mutations did not alter proximal-to-distal PAS ratio in CFIm68 KO cells. Normalized 300 nt/3.3k PAS ratio of mutated UGUA4+5 in *Pten* increased from 1.4- to 1.7-fold in CFIm59 KO over the parental line, while this ratio decreased near or at the level (~1.1-fold) of the WT reporter in CFIm68 KO (Figure 3C, 3E).

These data demonstrate the competitive regulation of proximal and distal PAS in *Pten* mRNA through UGUA sites. They are consistent with CFIm68 being the main direct regulator of distal (3.3k) UGUA elements 4 and 5, and with a partial compensation on proximal UGUAs 1-3 by CFIm59 upon CFIm68 loss. Lastly, these experiments further support distinct specificities for CFIm59 and CFIm68 on UGUA elements (See Discussion and Model in Figure 6).

### Distinct and opposing roles for CFIm59 and CFIm68 on global APA

We investigated the transcriptome-wide changes in APA upon impairment of CFIm complex subunits. 3’UTR-seq was performed on NIH3T3 cells following CFIm25, CFIm59, or CFIm68 KD, and the usage of distal and proximal PAS was quantified. Consistent with previous observations for *Pten* mRNA, CFIm25 and CFIm68 KD showed 10 to 12 times more downregulated distal PAS than upregulated, while there was 9 to 10 times more upregulated proximal PAS than downregulated (Figure 4A, top), thus supporting a transcriptome-wide shortening of 3’UTRs through APA. In contrast, CFIm59 KD resulted in 3.9 times more upregulated distal PAS than downregulated, and 2.6 times more downregulated proximal PAS than upregulated. CFIm59 KD resulted in a global lengthening of 3’UTR through APA (Figure 4A top). Similar analyses performed on the published PAS-seq of HEK293T CFIm59 KO and CFIm68 KO isogenic lines (19) led to comparable conclusions, although with fewer captured APA events (Figure 4A bottom), possibly due to clonal adaptations. Further analyzing these transcriptome datasets, a score for the Relative Expression Difference (RED) (32) of distal and proximal PAS usage was allocated for all CFIm-responsive gene passing quality control (see Materials and Methods). A positive RED score reflects an overall increase in distal over proximal PAS usage, whereas a negative RED score indicates the opposite, with an overall increase in proximal over distal PAS. KD of CFIm25 and CFIm68 in NIH3T3 resulted in negative RED scores for 77% and 75% of all responsive genes, respectively (Figure 4B, left). Similarly, in HEK293T, 85% of the responsive APA genes gave negative RED scores upon CFIm68 KO (Figure 4B, right). In comparison, 66% and 68% of CFIm-responsive genes produced positive RED scores upon CFIm59 KD in NIH3T3 and KO in HEK293T, respectively (Figure 4B). These results further support opposing functions for CFIm59 and CFIm68 in APA regulation at the transcriptome level. The current view is that CFIm recognition of UGUA elements is mediated through CFIm25 binding to RNA, and this subunit is a partner to both CFIm59 and CFIm68 subunits. Of the 5,934 genes that exhibit alternative 3’UTR isoforms in NIH3T3 cells, 80% (4,732) responded to at least one of CFIm subunit KD. When we queried the overlap of the 4,420 CFIm59 and CFIm68 responsive genes, only 32% (1,420) overlapped. Moreover, among the 3,939 genes that were oppositely responsive to CFIm59 or CFIm68 KD, only 22% (855) were shared, wherein CFIm59 KD favored distal PAS and CFIm68 KD promoted proximal PAS (Figure 4C). We noted that beside *PTEN*, many other well known genes involved in cancer followed a similar APA behavior. Among top hits along with *PTEN* were *BMPR2, ELAVL1*, and *NOTCH1* in HEK293T cells, as well as *Nras, Rictor*, and *Notch1* in NIH3T3 cells (Figure 4D, 4E). We further noticed several genes of the PI3K/Akt signaling cascade (see below).

**Figure 4.**
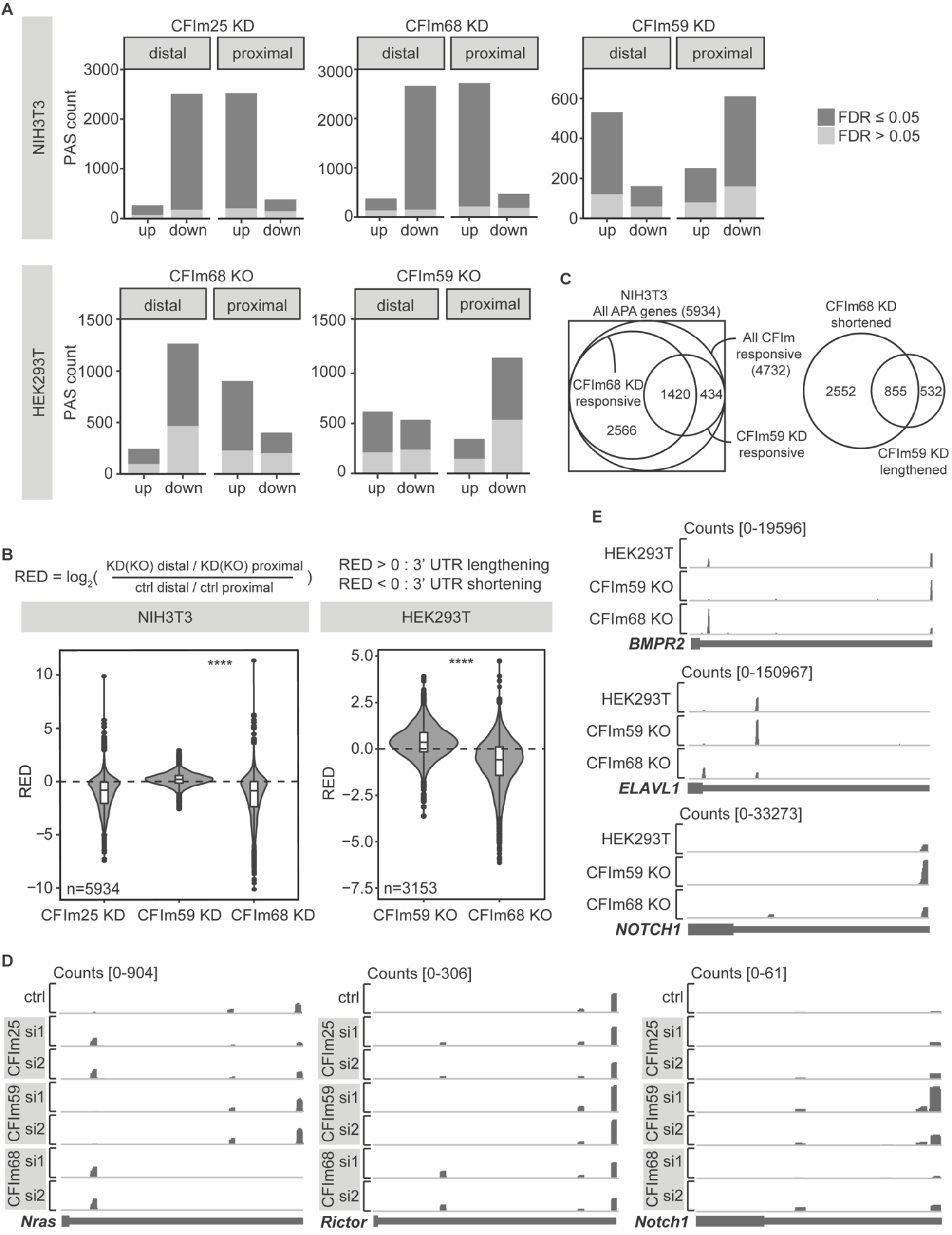
Distinct and opposing roles for CFIm59 and CFIm68 on global APA. (A) Up- and downregulated distal and proximal PAS counts upon CFIm KD or KO. Analyses of our NIH3T3 3’ mRNA-seq and public datasets of HEK293T PAS-seq were performed to tabulate each PAS expression direction upon CFIm disruption. Statistical significance was calculated with DEXSeq or Fisher’s exact test and corrected for multiple testing. Dark gray indicates PAS differential expression with false discovery rate (FDR) ≤ 0.05 and light gray for FDR > 0.05. (B) Relative expression difference (RED) score distribution of NIH3T3 and HEK293T upon CFIm disruption. RED of each gene was calculated as the log2 difference between KD (or KO) distal/proximal and control distal/proximal ratio. Paired two-tailed Student’s *t*-test was used (\*\*\*\**P* ≤ 0.0001). (C) Overlap of 3’UTRs shortened upon CFIm68 KD and lengthened upon CFIm59 KD in NIH3T3. (D and E) Genome tracks of example cancer-related genes that lengthen upon CFIm59 depletion and shorten upon CFIm25 and −68 depletion in NIH3T3 (D) and HEK293T (E) cells.

Taken together, these results further support the opposing functions for CFIm59 and CFIm68 in APA regulation as a transcriptome-wide phenomenon, and that CFIm59 and CFIm68 targets only partially overlap. This further prompts a reconsideration of how CFIm PAS specificity is achieved (see Discussion).

### CFIm APA regulation in the PI3K/Akt pathway

The PTEN phosphatase hydrolyzes phosphatidylinositol 3,4,5-triphosphate (PIP3) to phosphatidylinositol 4,5-bisphosphate (PIP2), and this activity antagonizes PI3K/Akt (protein kinase B) signaling. Given the global role of CFIm in APA regulation, the physiological implications of *PTEN* APA regulation cannot be inferred without integrating its impact on the PI3K/Akt cascade. Among the genes detected in our datasets, 359 genes were assigned to the PI3K/Akt pathway by Kyoto Encyclopedia of Genes and Genomes (KEGG) (33–35), of which 136 (38%) expressed alternative 3’UTR isoforms (APA), and 88 (25%) were responsive to CFIm perturbations (Figure 5A). Assignment to the Akt signalling cascade by Gene Ontology (GO) (36,37) led to comparable proportions. Of 206 detected genes assigned to the cascade, 76 (37%) underwent APA and 44 (21%) were responsive to CFIm perturbations (Figure 5A). PI3K/Akt cascade components that are responsive to CFIm include upstream activators such as receptor tyrosine kinases (RTK), the extracellular matrix (ECM), and G protein-coupled receptors (GPCR), as well as downstream targets of *Akt* such as *TSC1, Myc*, and *Bcl2* oncogenes (Figure 5B). Perhaps most strikingly, the catalytic p110α (*Pik3ca*) and regulatory p55γ (*Pik3r3*) subunits of PI3K, as well as *Akt3* responded to CFIm25, CFIm68 and CFIm59 KD similarly to *Pten:* CFIm25 and CFIm68 KD leading to 3’UTR shortening with negative RED scores, and CFIm59 KD leading to 3’UTR lengthening with positive RED scores (Figure 5C, 5D).

**Figure 5.**
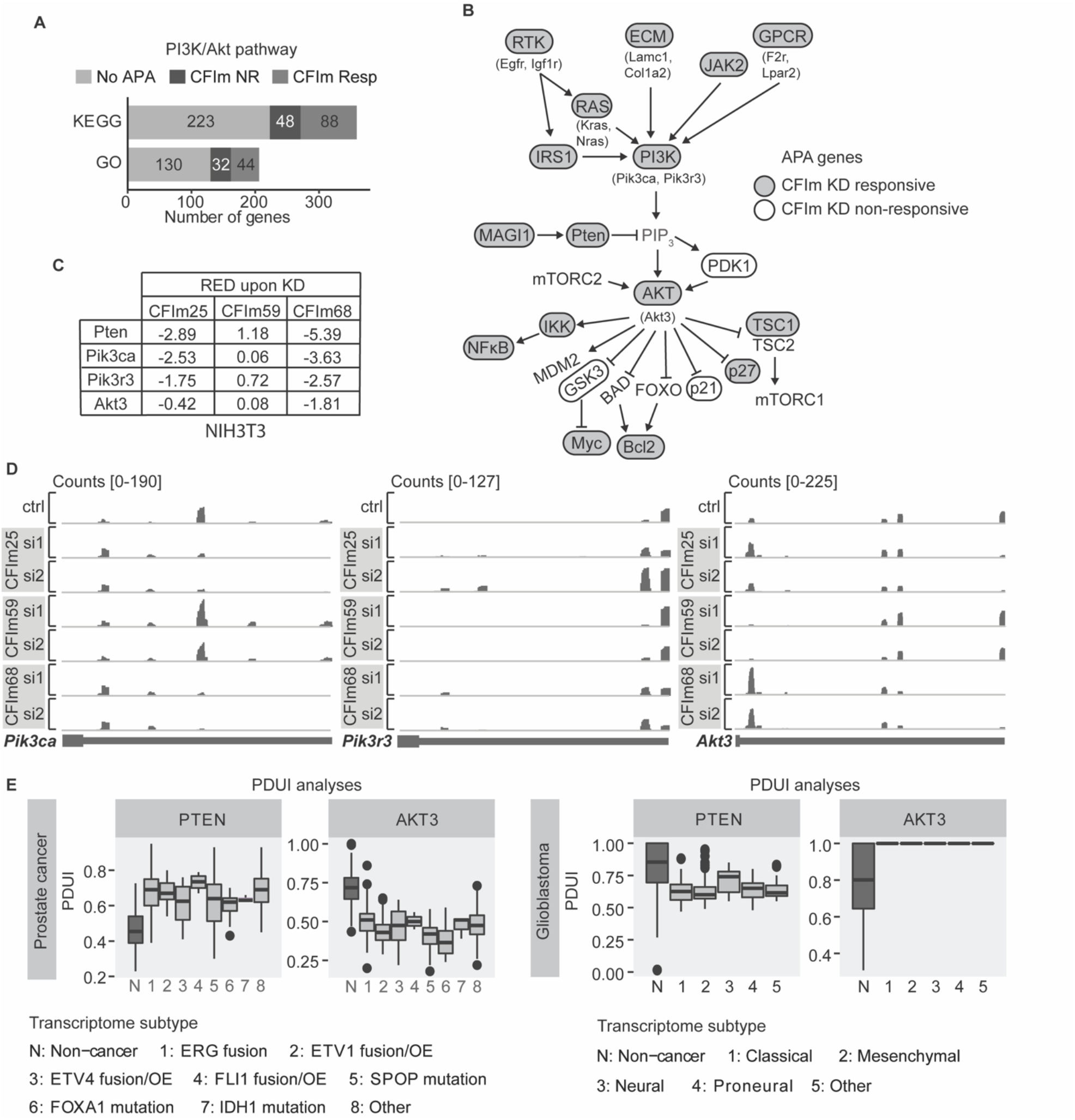
CFIm APA regulation in the PI3K/Akt pathway. (A) Tabulated APA status of genes designated to the PI3K/Akt pathway by KEGG, or the Akt signalling cascade by GO, from our NIH3T3 3’UTR-seq. No APA: genes with no detectable alterative 3’UTR isoform. CFIm NR: genes not responsive to CFIm KD. CFIm Resp: genes with detectable switch in 3’UTR isoform usage upon CFIm KD. (B) Schematic of the PI3K/Akt cascades. Genes responsive to CFIm KD are circled and shaded in gray, with example genes in brackets. Genes non-responsive to CFIm KD but with more than one detectable 3’UTR isoform are circled without shading. Genes with no detectable APA are not circled nor shaded. (C and D) RED scores (C) and gene tracks (D) of core PI3K/Akt components Pten, Pik3ca, Pik3r3, and Akt3, upon CFIm KD in NIH3T3. (E) PDUI analyses of PTEN and AKT3 in prostate cancer and glioblastoma. Each cancer is divided into transcriptomic subtypes defined by The Cancer Genome Atlas (TCGA) project as shown in the bottom legend. PDUIs of cancer datasets were collated from The Cancer 3’UTR Atlas (TC3A). PDUIs of non-cancer prostate and brain tissues were mined from APAatlas.

Pten and PI3K/Akt cascade misregulation is closely associated with several types of cancers. We next queried whether APA regulation of *PTEN* was altered in *PTEN*-driven cancers and compared it with other core PI3K/Akt cascade components. To enable enough depth across cancer subtypes, we relied on APA scoring by Percentage of Distal polyA site Usage Index (PDUI) from published mRNA-seq datasets. A higher PDUI means increased distal PAS usage (38). We integrated the PDUI scores of prostate cancer and glioblastoma patients of known subtypes from The Cancer 3’UTR Atlas (TC3A) datasets (38), with PDUI of normal tissues taken from APAatlas (39). In prostate cancers, and regardless of the transcriptomic subtype, the 3’UTR of *PTEN* was significantly lengthened. Curiously and in contrast, the 3’UTR of *AKT3* was significantly shortened, whereas no clear trend could be discerned from those datasets for PIK3CA and PIK3R3. In glioblastoma subtypes the trend was opposite, with *PTEN* mRNAs being expressed with more proximal PAS usage, and AKT3 3’UTRs being lengthened. Among other CFIm-responsive genes, PIK3CA 3’UTRs were significantly shortened as well (Figure 5E and Supplementary Figure 2). These results highlight the heterogeneity of APA changes across cancer subtypes and argue against onco-transcriptome 3’UTR misregulation through loss or impairment of any single CFIm subunit.

## DISCUSSION

Here we identify the CFIm complex as a direct regulator of *Pten* mRNA APA and show that this role has a significant impact on PTEN protein dosage. We further uncovered distinct and opposing roles for the CFIm59 and CFIm68 subunits in APA regulation on *Pten* mRNA and globally at the transcriptome level. CFIm59 depletion led to 3’UTR lengthening, while CFIm68 depletion led to 3’UTR shortening and favored distal PAS use. These results prompt a reconsideration of how CFIm specificity is established and reveal the breadth of APA regulation in the PI3K/Akt signaling cascade, as well as transcriptome-wide.

Even slight changes in PTEN dosage impact the output of the PI3K/Akt cascade, and its oncogenic down-regulation contributes to cancer initiation and progression (40). While CFIm perturbation results in an increase in both *PTEN* mRNA and protein expression, APA is merely one component of a much broader network of transcriptional, post-transcriptional, and post-translational regulation mechanisms controlling PTEN dosage (41–44). For example, part of *Pten* mRNA upregulation upon CFIm KD could be indirectly affected by *Pten* transcriptional regulators such as EGFR (45) and NFkB (46) which are CFIm sensitive. A recent bioRxiv prepublication manuscript identified *PTEN* transcriptional enhancers as one of the contributors to its APA regulation (47). Furthermore, another recent study proposed a mechanism of PTEN protein destabilization through APA regulation of WWP2 E3 ubiquitin ligase by CFIm59 (48). As such, the contribution of CFIm itself on PTEN dosage may not solely result from its direct involvement in *PTEN* mRNA APA.

How 3’UTR isoforms, individually or as co-expressed combinations, control or contribute to *PTEN* mRNA translation is also the result of input from a complex regulatory network. The 3.3k *PTEN* 3’UTR, by far the best studied of its mRNA isoforms, is the nexus of several post-transcriptional regulation mechanisms (41–44). Dozens of miRNAs are predicted to bind this 3’UTR, and its competition with a pseudogene served as conceptual framework for the ceRNA hypothesis. Even such indirect connections seem to exert influence on PTEN dosage; QTL analyses suggested that the *PTEN* 3’UTR pseudogene ceRNA drives a significant part of PTEN variation in cancers (49). It would stand to reason that longer 3’UTR isoforms, which encode more predicted regulatory elements, are better repressed by miRNAs, and that shorter isoforms are derepressed. However, our previous work instead revealed that longer 3’UTR isoforms contribute to the bulk of PTEN dosage, are largely resistant to miRNA regulation (presumably through folding structures (50)), and are in fact more stable than shortest isoforms (13). In agreement with this, a drastic 6- to 10-fold increase in the short 300 nt *Pten* 3’UTR isoform upon CFIm68 KD only resulted in a 1.5-fold increase in PTEN protein, and a similar increase was observed upon CFIm59 KD, with merely 2- to 4-fold increase in the long isoforms (Figures 1 & 2). In light of our results and given the multi-layered regulation converging on the 3’UTRs of *PTEN* mRNAs, a direct assessment of their translatability under physiological and oncogenic contexts appears as an important next research frontier.

### Distinct selectivity for CFIm59 and CFIm68 in APA regulation

Our results are consistent with a model whereby CFIm59 and CFIm68 have partially overlapping – but distinct – preferences for specific subsets of UGUA elements and functionally compete in directing PAS usage on pre-mRNAs that bear several UGUA clusters such as *PTEN* (Figure 6A). Consistent with this model, CFIm59 and CFIm68 can partially compensate for each other upon loss of one of the subunits (Figure 6B). This is illustrated by our analyses of UGUA mutations in wt and CFIm59- and CFIm68-KO cell lines (Figure 3). Indeed, CFIm68 appeared to drive most of UGUA4 and 5 input on the 3.3k isoform PAS, this activity was bolstered in CFIm59 KO, and UGUA4 and 5 still selectively promoted the nearby PAS in CFIm68 KO cells. Our model is consistent with prior reports of partial compensation between CFIm59 and CFIm68, namely in the recruitment of the CPSF subunit Fip1 to promote cleavage (19). Neither is it at odds with the reported enrichment of UGUA sites upstream of distal PAS, which had been regarded as a mechanism of distal PAS promotion by CFIm. However, that prior model did not account for the many cases such as *Pten* mRNA, where multiple UGUA sites are encoded near highly utilized proximal PAS, nor for the opposing functions of CFIm59 and CFIm68 subunits. CFIm-responsive transcripts that encode both proximal and distal UGUA elements identified in our analyses thus revealed a more complex dynamic between the CFIm subunits.

**Figure 6.**
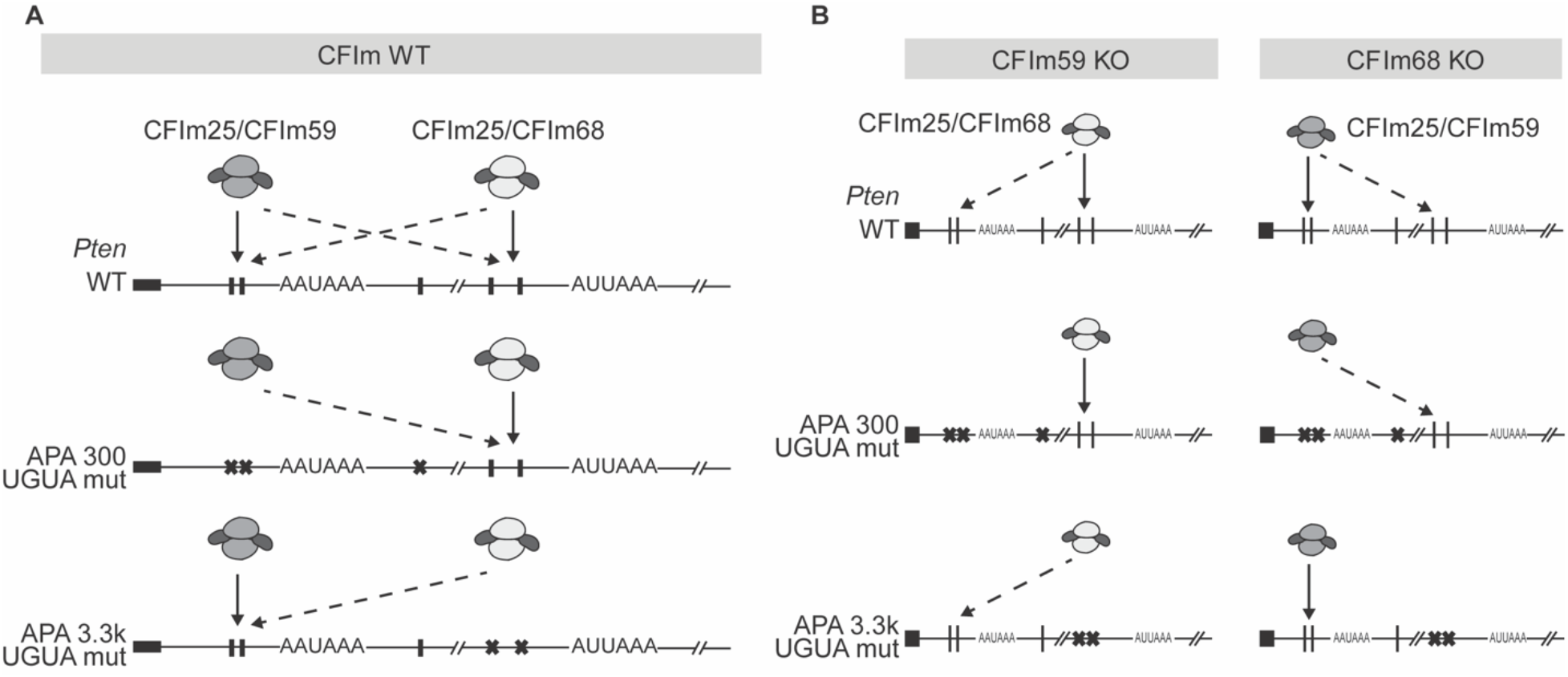
Model of *Pten* APA regulation by CFIm. CFIm25/CFIm59 complex promotes usage of APA 300 nt more efficiently (solid arrow) than APA 3.3k (dashed arrow), whereas CFIm25/CFIm68 is a better promoter of APA 3.3k than APA 300 nt. In the presence of both CFIm59 and −68 (A), loss of UGUA motif around either major PAS leads to APA shift in favor of the other PAS. In the absence of either CFIm59 or −68 (B) the same trend of APA shift occurs with UGUA mutations due to the partial compensation between the two complexes, but the magnitude depends on the targeting specificity of the complex present.

It may be hypothesized that CFIm59 acts as a negative regulator of CFIm68 through their direct interaction in a sub-population of CFIm complexes. This possibility could be considered as CFIm59 and CFIm68 can co-immunoprecipitate from HEK293 and glioblastoma cell lines (51,52). However, overexpression of CFIm59 does not mimic the effect of CFIm68 depletion on APA, at least for some of the CFIm-sensitive mRNAs (20). In light of our results, a more likely explanation is that distinct CFIm59 and CFIm68 CFIm complexes, associated with different UGUA elements, compete in their output on a subset of PAS. In line with this, only 32% of CFIm-sensitive PAS are responsive to both CFIm59 and CFIm68 KD. This is further supported by the coupled expression of the CFIm25 and −68 subunits, or CFIm25 and −59, while CFIm59 and −68 KD do not affect each other (Figure 1D).

The CFIm25 subunit directly binds to the UGUA elements in pre-mRNAs, but it is likely that CFIm25/CFIm59 achieve a distinct preference from CFIm25/68 complexes by directly contacting nearby additional sequences or through RNA-binding co-factors. CFIm59 and CFIm68 share an overall protein structure reminiscent of classic splicing SR proteins, which consist of a central proline-rich region flanked by an N-terminal RRM and a C-terminal RS-like domain. On their own, subtle structural differences in the width of the RNA exit cleft could lead to distinct RNA preferences (53). The most obvious difference between the CFIm68 and −59 subunits is the glycine-arginine rich (GAR) motif missing from CFIm59. Furthermore, the RS region of CFIm68 is thought to mediate interactions that are distinct from those of CFIm59 (54). Lastly, potential interactors of CFIm59 captured by co-fractionation and BioID RNA binding proteins including HuR and HNRNPA1, have previously been implicated in APA (55–59). Any of those differences may also contribute to the specificity of PAS enhancement.

### Breadth of APA regulation in the PI3K/AKT/PTEN axis and cancer

In our datasets, 80% of all genes identified with multiple functional PAS were responsive to depletion of at least one of the CFIm subunits. Several of these genes, including genes with opposing responses to CFIm59 and CFIm68 perturbations, are embedded in the PI3K/AKT/PTEN cascade. This consolidates CFIm as a master regulator of APA but also raises critical questions on the significance of this regulation in cancer. Considering the breadth of APA regulation on transcriptomes and their sensitivity to cellular and physiological changes, it should be expected that the positioning of regulatory elements in 3’UTRs has been harmonized through evolution for a fine-tuned output, whether enhancement or inhibition in the right cells and at the right time. In line with this concept, a previous 3’UTR-seq survey revealed the enrichment of miRNA-binding sites immediately upstream of APA sites in both pro-differentiation and anti-proliferative pre-mRNA sequences, which is expected to lead to a more potent output (60). As PI3K/AKT/PTEN signalling controls cellular states including proliferation, one could view the outcome of its signaling and the APA regulation among the mRNA of its core components as a feedback network for this crucial cascade.

Landmark papers have highlighted the prevalence of APA dysregulation in cancer (6,38,61). A persisting view that originates from those publications is that cancer cells tend to shorten 3’UTRs to ‘evade’ regulation by miRNAs and/or other trans-acting factors. Our APA analyses in tumor RNA-seq stratifications indicate that the outcome varies in different types of cancers, and loss of any individual CFIm subunit cannot account for the surveyed changes. This can be further inferred from several other publications. For example, CFIm25 perturbations were reported to contribute to glioblastoma tumor growth *in vivo* (24). A significant correlation was established between worst outcomes (poor survival) in glioblastoma patient and reduced CFIm25 and CFIm59 expression, but not with CFIm68 (51). In contrast, greater expression of CFIm59 and CFIm68 has been associated with increased tumor burden in mouse models of liver cancer (48,62). We reason that ultimately, the selective pressure for or against APA in any gene will very likely depend on the function of the gene itself, on the structure and sequence of its 3’UTRs, but also on the gene regulatory network that prevails in a specific cell fate and state. Consequently, the cellular impacts of *PTEN* APA should be considered in its cellular context and integrated with other APA regulation in the PI3K/Akt cascade. It may be premature to predict the oncogenic significance of any APA regulation or mis-regulation until comprehensive catalogs of dynamic PAS, of their regulatory elements and of the cellular cues they integrate are completed. Beyond the CFIm complex and its subunits, only a handful of APA regulators have been identified, and our study is among the first to detail how specific regulatory elements impact on individual 3’UTRs. Much of this important work on APA lies ahead.

## Data Availability

3’UTR sequencing datasets are available at GEO under the accession number GSE183704.

## Supplementary Data

Supplementary data are available at NAR online.

## Acknowledgements

We would like to apologize to authors whose directly related work may have not been cited in this manuscript. We thank Dr. Wei Li for his invaluable feedback on the manuscript, as well as Dr. Mathieu Flamand, and Dr. Hamed Najafabadi along with his student Gabrielle Perron for their initial guidance on the bioinformatics analyses.

## Funding

This work was supported by the Canadian Institutes of Health Research (CIHR) grant (MOP-123352) to T.F.D.; personal funding from Ms. Georgette Duchaine, McGill University Rosalind and Morris Goodman Cancer Institute Canderel studentship award, Fonds de recherche du Québec Santé (FRQS) Master’s training award, and CIHR Doctoral Research Award to H-W. T.

## Conflict of Interests

The authors declare no competing interests.

## Notes

### Competing Interest Statement

The authors have declared no competing interest.

