## Supplemental materials for "Distinct, opposing functions for CFIm59 and CFIm68 in mRNA alternative polyadenylation of *Pten* and in the PI3K/Akt signalling cascade"

**Supplementary Table 1: Oligos used**

| ID | Use | Sequence |
|---|---|---|
| TDO679 | *Pten* isoform-specific qRT-PCR: Reverse transcription | GCGAGCTCCGCGGCCGCGTTTTTTTTTTT |
| TDO3108 | *Pten* isoform-specific qRT-PCR: PCR1 forward primer for APA 3.3k | TATGACAGTATTCACGATTAGCC |
| TDO886 | *Pten* isoform-specific qRT-PCR: PCR1 forward primer for APA 300 nt; *Pten* ORF qRT-PCR forward primer | GCGTGCAGATAATGACAAGG |
| TDO680 | *Pten* isoform-specific qRT-PCR: PCR1 reverse primer | CCAGTGAGCAGAGTGACG |
| TDO2197 | *Pten* isoform-specific qRT-PCR: qPCR2 forward primer for APA 300 nt | TGGCAATAGGACATTGTGTCA |
| TDO3734 | *Pten* isoform-specific qRT-PCR: qPCR2 reverse primer for APA 300 nt | CAA GTG TCA AAA CCC TGT GG |
| TDO3764 | *Pten* isoform-specific qRT-PCR: qPCR2 forward primer for APA 3.3k | ACCTGCCAGCTCAAAAGTTC |
| TDO3765 | *Pten* isoform-specific qRT-PCR: qPCR2 reverse primer for APA 3.3k | TGCTGCACAGCACAAGAGTA |
| TDO1610 | qRT-PCR: *Hprt1* forward primer | AAGCTTGCTGGTGAAAAGGA |
| TDO1611 | qRT-PCR: *Hprt1* reverse primer | TTGCGCTCATCTTAGGCTTT |
| TDO887 | *Pten* ORF qRT-PCR reverse primer | TCTGGATTTGATGGCTCCTC |

| | | |
|---|---|---|
| TDO3768 | *Pten* APA 5-6k qRT-PCR forward primer | GCTCAGCAAATGCGTACCTA |
| TDO3769 | *Pten* APA 5-6k qRT-PCR reverse primer | ACAAGTCACAGAAGCACACA |
| TDO7301 | *Pten* APA 300 nt UGUA1 mutation forward primer | AAGGTTGAGAAGCTGTGTCATGTATATACC |
| TDO7340 | *Pten* APA 300 nt UGUA1 mutation reverse primer | TTTTTAACTGGACAACAAGTGTCAAA |
| TDO7302 | *Pten* APA 300 nt UGUA2 mutation forward primer | AGCTGTGTCAAGAATATACCTTTTTGTGTCAA |
| TDO7303 | *Pten* APA 300 nt UGUA2 mutation reverse primer | ACACAACCTTTTTTAACTGGACAAC |
| TDO7304 | *Pten* APA 300 nt UGUA3 mutation forward primer | GGCTGATGAGAATACGCAGGAGTT |
| TDO7305 | *Pten* APA 300 nt UGUA3 mutation reverse primer | TGTCTCCACTTTTTATAAAACTGGAAT |
| TDO7306 | *Pten* APA 3.3k UGUA4 mutation forward primer | GCTCTGTGAGAAAATGCTATGCACT |
| TDO7307 | *Pten* APA 3.3k UGUA4 mutation reverse primer | CACTGCTGCACAGCACAAGA |
| TDO7308 | *Pten* APA 3.3k UGUA5 mutation forward primer | AAATATGACGAGAACAGGATAATGCCTC |

| | | |
|---|---|---|
| TDO7309 | *Pten* APA 3.3k UGUA5 mutation reverse primer | GTGTATCCTCAGTGCATAGCAT |

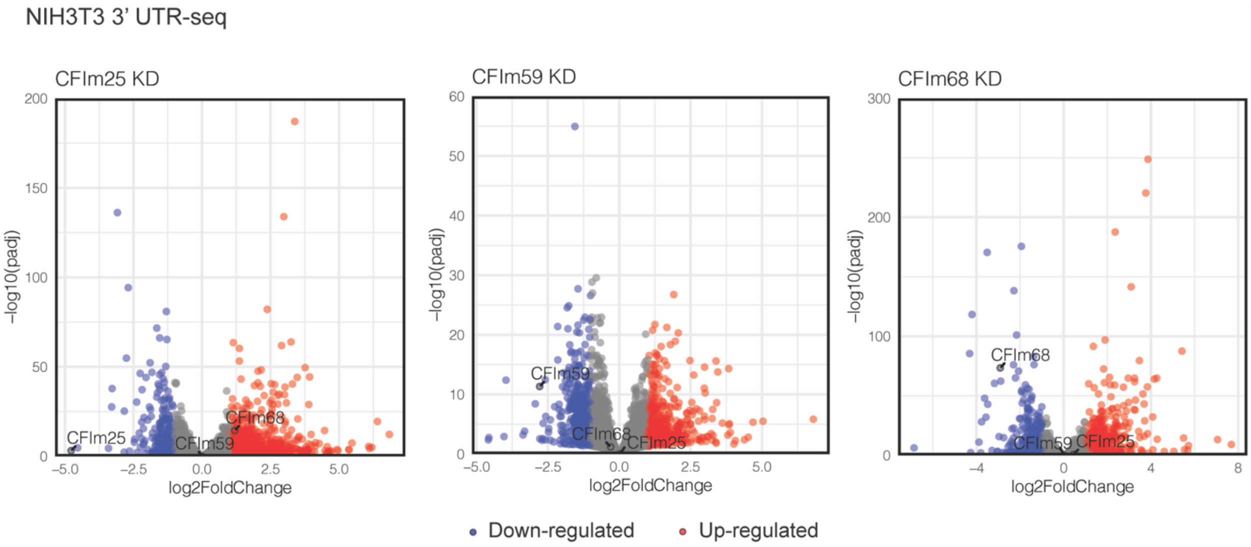


**Supplementary Figure 1.** Total RNA-seq analyses of NIH3T3 CFIm KD. Datasets of 3'UTR-seq were analyzed as total RNA-seq datasets by collapsing all mRNA isoforms of each gene and tabulating the total count. Up-regulated and down-regulated genes are shown as red and blue points, with CFIm components labeled. Fold change cut-off was set at 2-fold with false discovery rate adjusted P-value (padj) threshold of 0.05.

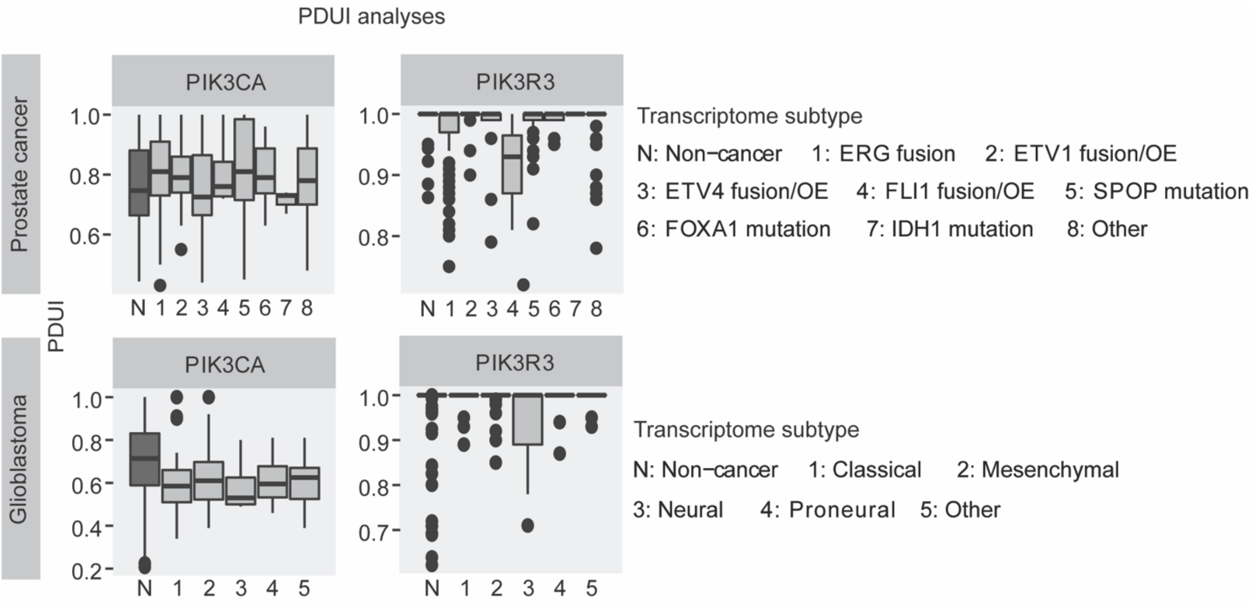


**Supplementary Figure 2.** PDUI analyses of PI3K subunits in cancer. APA changes of PIK3CA and PIK3R3 in prostate cancer and glioblastoma are profiled. Cancer PDUI datasets are taken from TC3A and normal tissue PDUI taken from APAatlas. Transcriptomic subtypes of each cancer are defined by the respective TCGA projects.
